# KASP-Hedgehog: Efficient and routine tool for allele specific primer design using multidisciplinary data types

**DOI:** 10.1101/522045

**Authors:** M.A Alsamman, M Abdelsattar, S.D. Ibrahim, Hamwieh Aladdin

## Abstract

The Integration of known nucleotide variations in breeding programs and medical assessments demands the ability to scan and validate thousands of gene-affected SNPs. Designing PCR primers targeting these SNPs require computational tools could generate accurate and target-specific primers using multidisciplinary data types with high accuracy. KASP-Hedgehog is a local-installation, simple and routine tool for allele-specific PCR primer design. KASP-Hedgehog gives user the ability to design KASP primer from a variety of genomic data, extract bi-allelic SNPs for genomic sequences with unknown nucleotide variations and select SNPs with potential effect depending on gene annotation. This tool can use user-provided or self-produced SNPs database to create allele-specific primers with degenerate structure in order to enhance PCR assay selectivity and increase amplification reaction. Additionally, it run an *in silico* PCR test for designed primers against provided FASTA sequences in order to increase primers selectivity and provide user more information. Moreover, it has interactive and user-friendly graphical user interface (GUI) in addition to a command-line package could be integrated in different bioinformatics pipelines.

## Introduction

Since the first published full genome study (1), the genome-sequencing avenue for different living organisms has expanded exponentially. The National Center for Biotechnology Information (NCBI) genome repository now holds more than 35 thousand organisms. Additionally, several plants, animal and microbial pan genomes or SNP databases are available for public use (2–4).

Simultaneously, molecular marker techniques have been advancing over time; it provides technologies deliver extremely high levels of assay robustness and accuracy with significant cost saving. An example of these technologies, KASP assay that utilizes a unique form of competitive allele-specific polymerase chain reaction (PCR) for the identification of genetic variations occurring at the nucleotide level to detect single-nucleotide polymorphism (SNPs) or InDels (5)□. KASP has been successfully developed and validated nucleotide variations related to important traits across different organisms (6–8) □.

The first step in integrating known nucleotide variations in breeding programs and medical assessments demands the ability to scan and validate thousands of gene-affected SNPs using high throughput allele-specific technologies such as KASP assay. Once these SNPs effects are validated, it could be integrated in disease detection, breeding programs or gene-editing assay.

The previous steps require vital computational tools could be used to design allele-specific PCR primers with high selectivity and efficiency. These tools must offer robustness, accuracy, the ability to test primers specificity through *in silico* PCR analysis and it could use the available information such as genome annotation, in order to select SNPs with potential. Moreover, the availability of SNPs variations retrieved using NGS technologies such pan genomes or simple sequence alignment could be used to create PCR primer with degenerate structure. This degenerate structure will add more selectivity to the PCR assay and extend the *in silico* PCR analysis performance and genomic regions with high mutation rate.

In the past few years, some bioinformatics tools have been introduced with the ability to design allele-specific PCR primers. WASP is a web-based allele-specific PCR assay-designing tool for detecting SNPs and mutations (9)□, it has the ability to design KASP primers using user-provided sequences or database SNP identity number. WASP has an option to perform *in silico* PCR against human genome, in order to filter primers depending on specificity. Prim-SNPing is an online tool provides interactive, user-friendly and cost-effective primer design for SNP genotyping. Prim-SNPing can accept multiple sequence formats for human, mouse, and rat SNP sequences as an input (10). These online tools cannot handle degenerate sequences (has more than one SNP), the *in silico* option PCR is only available for few selected genomes and there is no local installation version for these tools until now.

In this study we introduce KASP-Hedgehog as a simple and routine tool for allele-specific PCR primer design. KASP-Hedgehog is a local-installation tool gives user the ability to design KASP primer from a variety of genomic data. In addition to designing primers for know SNPs Hedgehog offers the ability to perform multiple sequence alignment, in order to extract bi-allelic SNPs for genomic sequences with unknown nucleotide variations and users can select SNPs with potential effect depending on gene annotation. Hedgehog can use user-provided or Hedgehog-produced SNPs database to create allele-specific primers with degenerate structure in order to enhance PCR assay selectivity and increase amplification reaction. KASP-Hedgehog run *in silico* PCR for designed primers against provided FASTA sequences in order to increase primers selectivity and provide user more information. KASP-hedgehog has interactive and user-friendly graphical user interface (GUI) in addition to a command-line could be integrated in different bioinformatics pipelines.

## Methodology

KASP-Hedgehog is local-installation software written using PERL (v5.18.2) and C programming languages. KASP-Hedgehog has a simple GUI and can be installed on LINUX and Microsoft Windows operating systems (**Figure 1**).

**Figure 1:**
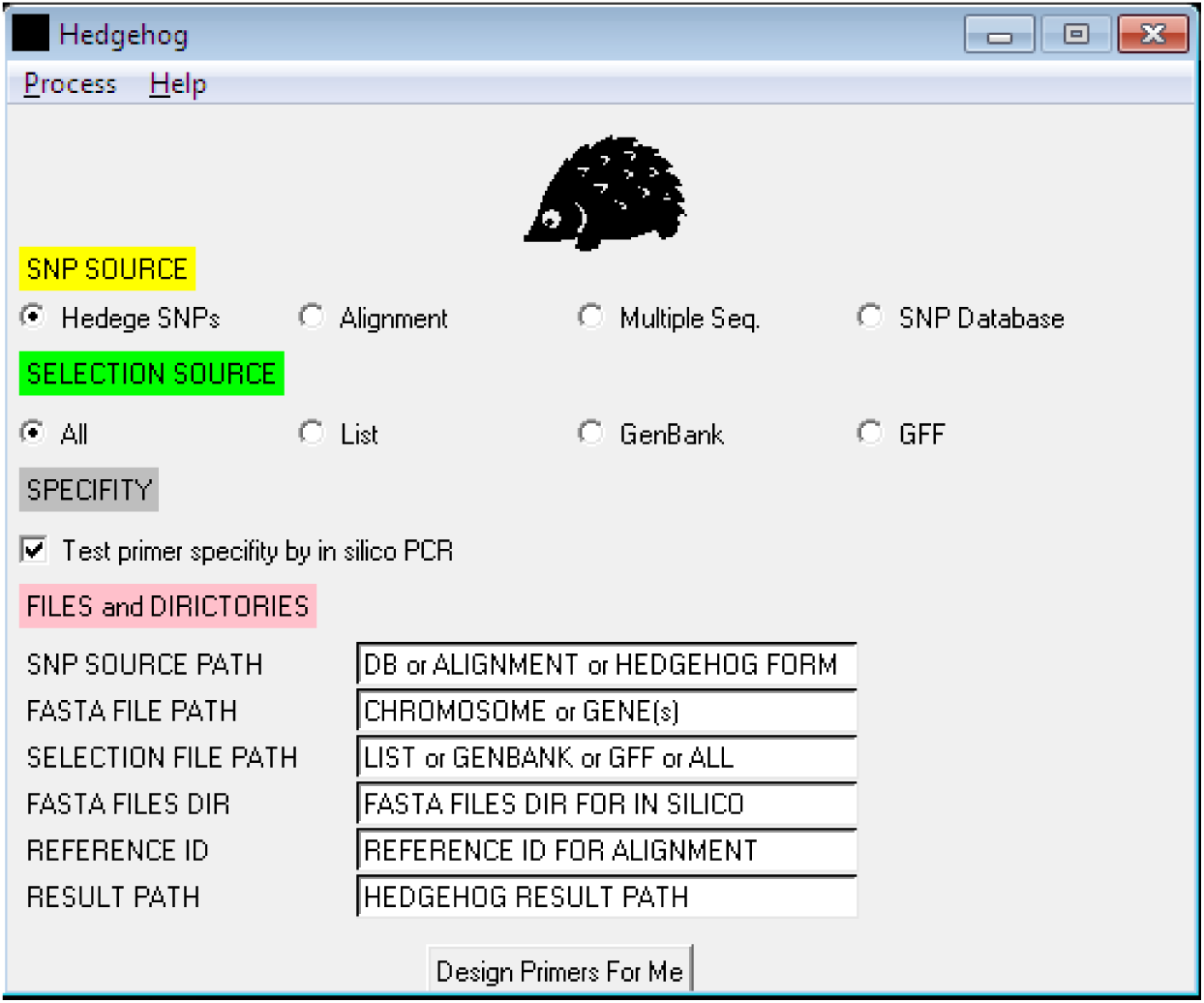
KASP-Hedgehog graphical user interface.

## Input

In order to give the user a multiple choices, KASP-hedgehog can handle different biological data such as a comma separated file contains different nucleotide sequences (could be degenerated according to IUPAC), where the targeted nucleotides are highlighted by “[“ and “[“ (Hedgehog format), multiple sequence file contains different sequences for the same gene (FASTA or CLUSTALW formats) or SNP database (tab delimited format). If user provided data in multiple sequence formats Hedgehog will align these sequences using MUSCLE (11)□ and use it to create a consensus sequence and SNP bi-allelic database for primer design. User can select specific SNPs located in interested genomic areas by providing a tab-delimited list, GFF or GenBank annotation files. Hedgehog will use all these information to create a multiple sequence file contain all possible nucleotide variants. These sequences will be used to design allele specific primers to target interested SNPs using PRIMER3. The output primers will be returned to PRIMER3 (12)□ if the sequences contain degenerate nucleotides in order to recheck its accuracy. The resulted primers will be used for *in silico* PCR analysis against user-provided sequences (could be whole chromosomes or multiple genes) (**Figure 2**).

**Figure 2:**
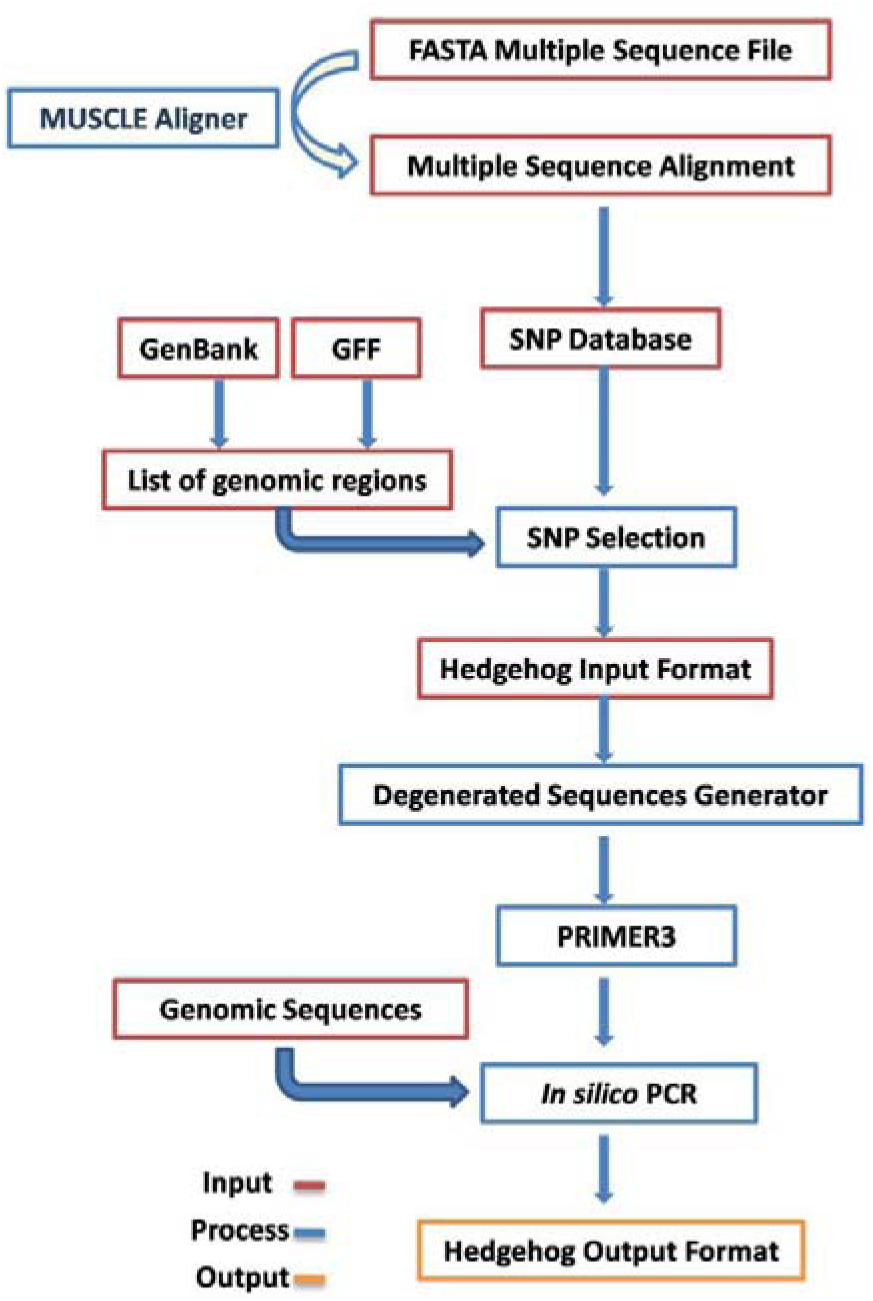
KASP-Hedgehog computational flowchart.

## Output

If successfully executed, KASP-Hedgehog will generate allele-specific primers, where two left primers and one common right primer target specific SNP. the tab delimited file will contain specific information regarding to these primer specifications such as SNP identity number, primers sequences, genomic positions, primer isoforms count (if these primers have degenerate nucleotides), number of isoforms failed to full-fill user-defined parameters, design penalty, melting temperature, GC content percentage, self-annealing value and other information. Additionally, *in silico* PCR a primers result, where amplification hits in the genomic sequences nucleotide has mismatches with nucleotide before user-defined 3’ nucleotide or after 3’ nucleotides are informed.

## Validation

Developing allele-specific SNP primers for plant genomes could used in genome selection and SNP genotyping is a main goal in plant breeding. KASP-Hedgehog was used to develop an SNP allele-specific primer database for chickpea. The published chickpea genome (13)□ and the SNP database (4)□ was used design 9811 allele-specific primers using default parameters, these primers target 2498 SNPs, which located in 540 genes.

The NCBI-PopSet is a database of DNA sequences which have been collected to study the relatedness within a particular population (www.ncbi.nlm.nih.gov/popset). Nowdays, the PopSet database contains 24077 human gene set, these genes was sequenced in order to study population,phylogenetic and mutation studies. Additionally, the Popset database could provide a useful source for nucleotide variations, converting these variations into high-throughput genotyping assays will provide major SNP resource for the dissection of phenotypic variations. KASP-Hedgehog was used to design 439 and 1188 allele-specific primers for 27 and 12 human and wheat genes, respectively. These primers was designed where one SNP could be targeted with 1, 2 or 3 different primers. Some of the designed primers have degeneracy and all primers have failed primers design specifications have been removed. The primers sequences and targeted genes are shown in **Supplementary 1.**

## Result and discussion

KASP-Hedgehog is local-installed software, where user can use to design more accurate allele specific primers using minimum information. Designing allele specific primers via multiple sequence alignment can be used to screen SNPs in species has no full genomes yet or has novel SNPs depending on the self created SNP database. Selecting SNPs using genome annotation or user-provided genic list could provide a cost-effective primers set for SNP genotyping in order to minimize genomic screened area dramatically. KASP-Hedgehog can handle degenerate nucleotides inside targeted sequences and can use them as a guide to design primers with as minimum as paring mismatches. Additionally, these nucleotides will give users another level of result preferences, where primers with high mis-pairing in their different primer degeneracy isoforms can be neglected or selected depending on depending on research target. The *in silico* PCR analysis integrated in KASP-Hedgehog procedure will give the user the ability to test primers selectivity. In addition, primers degeneracy will be exported to *in silico* to show all untended pairing with random genomic positions, in order to give users high form of primer specificity.

KASP-Hedgehog has a simple graphical user interface (GUI) and can be installed on different operating systems. This local version will give users the ability to handle new genomic sequences and SNPs and its programming scripts could be easily integrated in different bioinformatics pipelines or platforms.

## Availability

The open source codes, sample input and output data, Linux and Microsoft Windows installations are freely available at https://github.com/AlsammanAlsamman/KASP-Hedgehog. In order to run through command line or LINUX, KASP-Hedgehog uses PERL (v5.18.2). PERL-TK module is needed to run GUI.

## Author Contributions

A.M.A.; software development, implementation of the computer code and supporting algorithms and testing of existing code components. A.M. and I.S.D.: writing – original draft, validation and resources. H.A.; Development and design of methodology and supervision.

